# Quantitative imaging of intracellular density with ratiometric stimulated Raman scattering microscopy

**DOI:** 10.1101/2021.06.13.448254

**Authors:** Benjamin Figueroa, Fiona Xi Xu, Ruoqian Hu, Shuaiqian Men, Dan Fu

## Abstract

Cell size and density impact a wide range of physiological functions, including tissue homeostasis, growth regulation, and osmoregulation. Both are tightly regulated in mammalian cells. In comparison, density variation of a given cell type is much smaller than cell size, indicating that maintenance of cell type-specific density is important for cell function. Despite this importance, little is known about how cell density affects cell function and how it is controlled. Current tools for intracellular cell density measurements are limited either to suspended cells or cells growing on 2D substrates, neither of which recapitulate the physiology of single cells in intact tissue. While optical measurements have the potential to measure cell density *in situ* and noninvasively, light scattering in multicellular systems prevents direct quantification. Here, we introduce an intracellular density imaging technique based on ratiometric stimulated Raman scattering microscopy (rSRS). It quantifies intracellular drymass density through vibrational imaging of macromolecules. Moreover, water is used as an internal standard to correct for aberration and light scattering. We demonstrate real-time measurement of intracellular density quantification and show that density is tightly regulated across different cell types and can be used to differentiate cell types as well as cell states. We further demonstrate dynamic imaging of density change in response to osmotic challenge as well as intracellular density imaging of a 3D tumor spheroid. Our technique has the potential for imaging intracellular density in intact tissue and understanding density regulation and its role in tissue homeostasis.

## Introduction

Biological cells have widely varying size. However, cells from a given type often exhibit remarkable regularity in size, where abnormal size can be an indicator of neoplastic growth. Maintaining proper cell size is essential for a cell’s function in a multicellular organism.^1–3^ Size is measured as either mass or volume, and the ratio of these two parameters is mass density, which is reflective of macromolecular concentration and crowding that affects gene activity, biochemical reactions, and biophysical properties of the cell.^4–7^ Cell-type specific density has been commonly used in density gradient centrifugation for cell separation.^8,9^ Recent single cell measurements show that intracellular density is significantly more closely regulated than cellular mass and volume.^10,11^ Interestingly, under normal growth conditions, mammalian cells density is remarkably constant throughout the cell cycle except during mitosis,^12,13^ but intracellular density changes in response to a number of physiological and pathological conditions. For example, lymphocytes density decreases when activated and increases when becoming memory cells.^14,15^ The tight regulation of cell density suggests that it not only can be used to identify cell types but also reveal changes in cell states of single cells.^7,11,16^ Despite its importance, we know little about which cellular functions are affected by intracellular density, how density influences cellular processes, and how density is regulated in response to physiological conditions.

One main obstacle to study density regulation and its function is accurate and noninvasive measurements of density at the single cell level. Although density gradient centrifugation can isolate cells of a particular density, it requires suspended cells, and the centrifugal force-induced stress may affect cell survival. It is also not suitable for single cell analysis. More recently, suspended microchannel resonators (SMRs) have been developed to precisely measure the buoyant mass of single cells.^17^ By weighing single cells in solutions at two different densities, density of single cells can be determined.^10^ However, SMR measurements are restricted to suspended cells with limited throughput. It has no spatial resolution and thus cannot resolve cellular organelles.

Cell density can also be indirectly measured with noninvasive optical imaging methods. The main technique is based on quantitative phase microscopy (QPM, also known as digital holographic microscopy).^18^ QPM is a technique that measures the optical phase delay induced by the refractive index difference between a sample and medium. The extent of this phase delay is an integral of sample thickness and its average refractive index (RI). Because RI is linearly proportional to cell drymass density (i.e. mass density of all biomolecules, mostly including proteins, lipids, nucleic acids, and carbohydrates)^19^, QPM directly measures cell drymass.^20^ To obtain intracellular density, quantitative phase tomography (QPT, also known as optical diffraction tomography) computationally reconstructs the RI maps in 3D from multiple QPM images at different illumination angles.^21–23^ Compared to SMR, QPT works with either adherent or floating cells and offers submicron spatial resolution. For example, QPT showed that nucleus is a less dense organelle compared to the rest of the cell, unlike previously thought.^24,25^ However, the need for 3D tomographic measurement with high numerical aperture lens and computational reconstruction limits the throughput of QPT. Moreover, QPT uses single scattered light and it is difficult to work with thick samples that have multiple scattering.^26^ There has been recent progress in overcoming this challenge, but the RI accuracy in tissue imaging remains to be determined.^27,28^

Here we report an intracellular density imaging by ratiometric stimulated Raman scattering (iDI-rSRS) technique that overcomes the limitations of existing techniques. SRS microscopy is an powerful chemical imaging technique which quantifies the spatial distribution of molecules based on their intrinsic vibrational contrasts.^29,30^ Because SRS intensity is linearly proportional to molecular concentration, SRS has been widely used to measure endogenous and exogenous concentrations of small molecules in living cells.^31,32^ SRS at C-H stretching frequency (or C-D, in the case of deuterium labelling) and spectral unmixing has also been used to qualitatively determine the relative content of major macromolecules including proteins, lipids, and nucleic acids.^33–35^ Recognizing that protein accounts for 60% of the drymass of a mammalian cell and the relative composition of biomolecules in cells are stable,^36^ it is possible to use C-H SRS imaging at the protein peak to probe cell drymass density. In this manuscript, we demonstrate the first reported quantitative intracellular cell density imaging with SRS. To calibrate contribution of other macromolecules to cell drymass, we directly compare SRS images of total protein mass to QPM measured drymass. The obtained calibration factor allows us to directly convert SRS intensity to drymass density. To overcome signal loss due to the ubiquitous presence of sample induced aberration and light scattering,^37,38^ especially in multicellular samples, we developed a novel ratiometric SRS method to normalize protein SRS with water SRS and compensate these effects. Our iDI-rSRS technique provides facile and real-time intracellular density mapping of living cells in both 2D and 3D at submicron resolution. We demonstrate real-time measurement of intracellular density quantification and show that density is tightly regulated across different cell types. Variations in density can be used to separate cell types or cell states such as mitosis and apoptosis. Dynamic imaging of intracellular density reveals osmotic response of cells. We further show that cells grown in 3D environment has higher intracellular density than those grown in monolayer culture. This advancement paves the way for real-time label-free *in vitro* and *in vivo* 3D imaging of single-cell mass density and serves to address many open biological questions in density regulation, macromolecular crowding, and their functional importance in cell physiology and tissue homeostasis.

## Methods

### Cell culture

A549, HeLa, and HEK293 (ATCC) were maintained at 37°C in an atmosphere of 5% CO_2_ (vol/vol) humidified incubator. A549 is cultured in Ham’s F-12K medium (Gibco, Catalog number: 21127022). HeLa and HEK293 were cultured in Dulbecco’s Modified Eagle Medium (DMEM) (Gibco, Catalog number: 11885084). All culture media were supplemented with 10% fetal bovine serum and 1% penicillin-streptomycin. For both fixed and live monolayer cells imaging, cells were seeded on coverslips 48 hours before the imaging session. Experiments implementing static cells were fixed with 4% paraformaldehyde (15 mins) at room temperature. The fluorescent dyes used were Hoechst 33342 and CF®488A Annexin V to stain the nucleus and apoptotic cells, respectively. All dye protocols were based on the provided instructions from the manufacturer.

To grow 3D spheroids, HEK293 cells were cultured in AggreWell™ 24 well-plates 48 hours before imaging. Each well was pre-treated with anti-adherence rinsing solution before seeding the cells. The seeding density for HEK293 per well was achieved by taking 1 ml of single cell suspension of approximately 6*10^4^ cell/mL and 1 ml of complete culture medium with estimation of 50 cells per microwell (final volume of 2 mL per well). The spheroids were incubated for one hour before imaging.

### Broadband rea-time ratiometric SRS imaging

A femtosecond dual-beam laser system (Insight DS + from Spectra-Physics) was used for hyperspectral SRS imaging based on an orthogonal modulation spectral focusing scheme as described in our earlier publication.^39^ Briefly, we used a parabolic fiber amplifier to increase the Stokes laser bandwidth from 6 nm to ∼40 nm,^40^ increasing the bandwidth from the original laser system of ∼200 cm to ∼500 cm^-1^, which allows us to acquire both C-H region at 2935 cm^-1^ and the lower shoulder of the Raman spectral signature of water around 3190 cm^-1^. We then employed an orthogonal modulation technique on our Stokes laser for simultaneous dual-band imaging, which was previously used for real-time microscale thermal imaging with SRS.^39^ The two orthogonal polarization pulse trains were then passed through a birefringent crystal to provide the temporal delay needed to probe two desired SRS transitions for normalized mass density imaging. Simultaneous two-channel SRS imaging is important to mitigate measurement errors from laser intensity fluctuation and motion artifact. The pump beam was chirped by 30 cm of high dispersion H-ZF52A glass rods to match the chirp of the Stokes beam. The combined beams were directed into a home-built inverted laser scanning microscope (Olympus IX73). A 60x oil immersion objective (Olympus UPLXAPO60XO) was used for imaging. Two SRS images were acquired simultaneously from the lock-in X and Y output (Zurich Instruments HF2LI) at a frame (512 x 512 pixels) rate of 0.5 frame/sec unless specified otherwise. Power of pump and Stokes were set at 40 and 45 mW, respectively unless specified otherwise.

### SRS calibration and cell imaging

Calibration of SRS wavenumbers was done by imaging the Raman peaks of dimethyl sulfoxide (DMSO) and methanol and performing linear fitting (Fig. S1). To measure mass density, we simultaneously image the CH_3_ mode pertaining to proteins (∼2930 cm^-1^) and the OH mode of water (∼3170 cm^-1^) as indicated in the shaded region in Fig. 1A. The normalized mass density distribution of the imaging plane is calculated from the intensity ratio (I_Ratio_) of the two SRS images (I_2935 cm_^-1^/I_3190 cm_^-1^) (for details, see supporting information: drymass calibration). For calibration, we measured I_Ratio_ as a function of protein concentration. As expected, I_Ratio_ scales linearly with concentration (Fig. 1B). The calibration can then be used to convert I_Ratio_ to protein density. From the measured SRS intensity ratio error, we determine that SRS can provide quantitative density measurements with an accuracy of 0.7 fg/μm^3^. To image live cells, a temperature-controlled micro-observation chamber was added to our microscope (FCS2, Bioptechs) which was kept at a constant temperature of 35 °C. The FCS2 uses an electrically conductive transparent thin film of indium-tin oxide to provide uniform heating. In addition, we used a Bioptechs Objective Heater System to eliminate the thermal heatsink from the objective.

**Figure 1:**
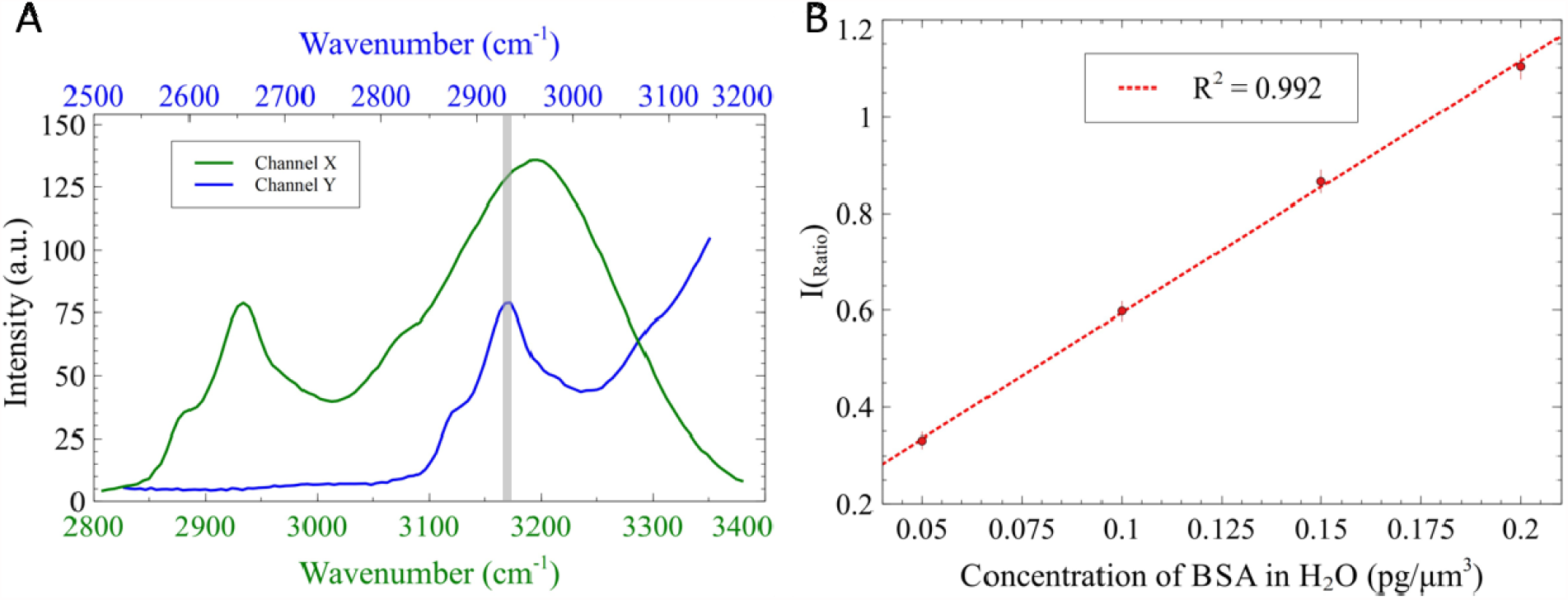
A) Simultaneous collection (shaded region) of 2-channel SRS spectra arising from CH and OH stretching vibration of 10% BSA in H_2_O. B) Calibration showing the linear relationship between protein mass and SRS I_Ratio_. The standard deviation of the intensity ratio for each image is indicated as the error bar.

### QPM image acquisition and drymass calculation

The QPM images were collected with a SID4-HR GE sensor (Phasics) mounted to a Nikon Eclipse TE2000-E microscope.^41^ A Nikon CFI Plan Apo Lambda 20X objective was used for imaging. The acquisition and processing of the phase images were accomplished with the SID4 Bio software. The phase images were segmented with the ROI (region of interest) toolbox in the SID4 Bio software. The cell optical volumes were determined by integrating optical phase over the area of the cells. The cell drymasses were then calculated by dividing the cell optical volumes by a conversion factor, 0.18 µm^3^/pg, known as the specific refractive index increment.^19^

### Image segmentation

We first convert the rSRS intensity into mass density using the calibration of BSA in water. Since SRS provides mass density measurement in 3D, total drymass can be determined by integrating the density over the segmented volume of a cell. For SRS images, the first step in the segmentation process is to determine the threshold value used to evaluate boundary pixels containing the cell’s volume. This was done by finding the FWHM value of the cell’s intensity and thresholding the lower value pixels out. The boundary is used to calculate the volume of each segmented cell. At the boundary, the cell SRS signal will be convoluted with signal from outside the cell due to the limited axial resolution at ∼0.9 μm. To reduce uncertainty in mass quantification, we padded our volume masks with 1 pixel layer to obtain pixels used in integrating cellular mass.

### Osmolarity challenge experiments

For osmolarity challenge experiments, cells are grown as monolayer in a temperature-controlled on-stage cell culture chamber (FCS2 from Bioptechs). The growth media contained in the FCS2 chamber was exchanged at a flow rate of 50□μl□min^−1^ using a Harvard Apparatus syringe pump. For hypotonic exchanges, F12K medium (300<u>□</u>mosM□kg^−1^) was diluted with water to 150□mosM□kg^−1^. In the case of hypertonic exchanges, F12K was supplemented with 5 mM NaCl to increase osmolarity to 500□mosM□kg^−1^.

## Results and Discussion

To calibrate rSRS measured drymass, we compared our rSRS measurements with QPM (Figure 2A-B). Mass of SRS imaged cells are obtained by converting rSRS to protein density using calibration in Figure 1B and then integrating protein density over the 3D cell volume. Using three different cell lines we obtained strong linear correlation (.99) between rSRS mass and QPM mass with a RMSE of 42 pg (Fig 2C). The slope is the correction factor *γ*, which is 1.018. The drymass density of a cell can then be easily determined by simply multiplying rSRS mass density with *γ*, which accounts for the variations in SRS intensity of different biomolecules (see supporting information). It is important to note that the correlation is degraded due to segmentation errors in 3D for SRS. The drymass density is thus more accurate than integrated mass measurements.

**Figure 2:**
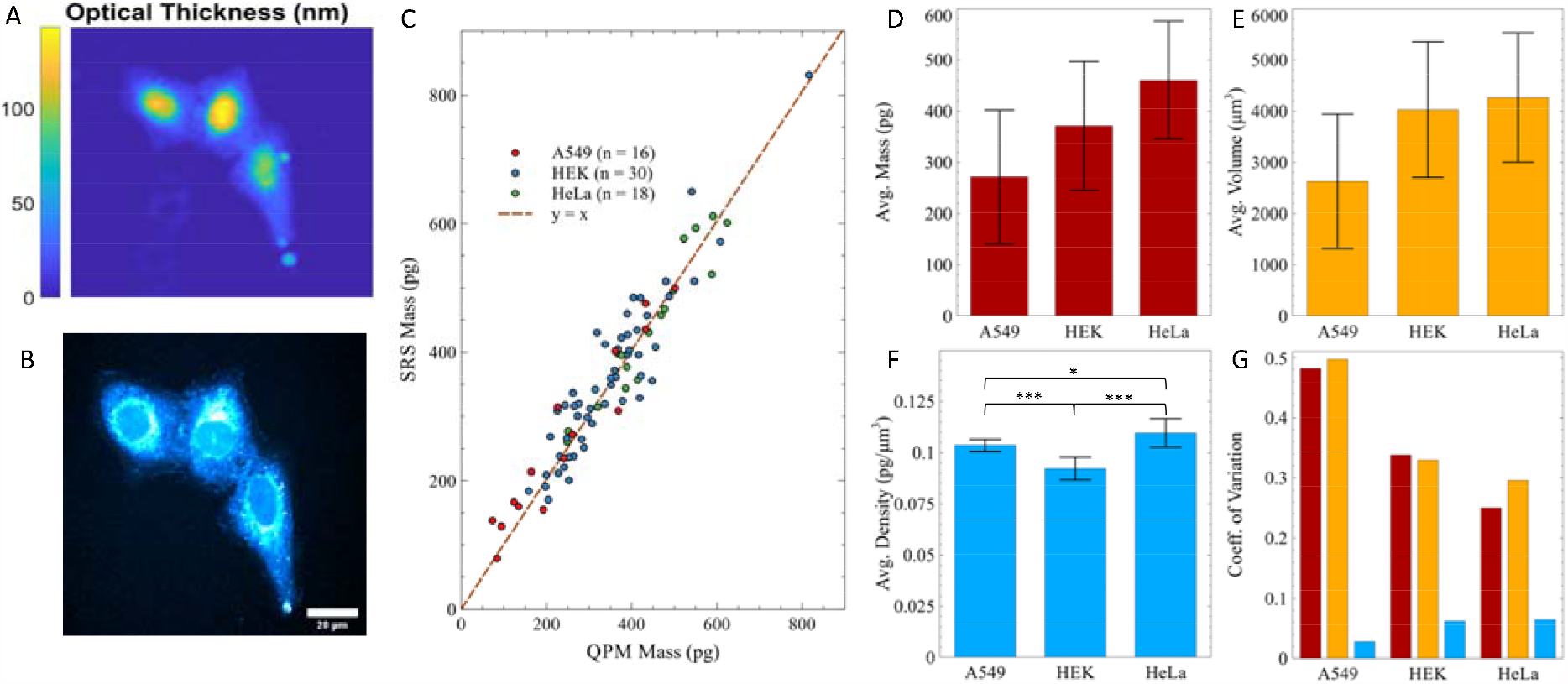
Validation of SRS microscopy to accurately measure physical parameters. Representative A) QPM and B) SRS image of A549 cells. C) Correlation between QPM mass measurements and SRS mass measurements for three different mammalian cell types. Average D) mass, E) volume, and F) density between the three different cell types. G) Comparison of the coefficient of variation between mass, volume, and density. Error bars are ± one standard deviation of the mean. *p≤0.05, *** p≤0.001

From our measurements, we observed that the HeLa cells have a higher average mass (460 ± 115 pg) than A549 (271 ± 130 pg) and HEK293 (371 ± 125 pg) cells, with the large standard deviations resulting from cells with different sizes and stages within the cell cycle (Fig. 2D). Similarly, we observe large variation in cellular volume (Fig. 2E). Consistent with previous observations, whereas cellular mass and volume can vary by as much as 50%, density is tightly regulated.^10^ From Fig. 2F, we observe that HEK293 cells have the lowest density (0.092 ± 0.0057 pg/μm^3^), followed by A549 cells (0.103 ± 0.0029 pg/μm^3^), and HeLa cells (0.109 ± 0.0071 pg/μm^3^). These values also agree with previous measurements using QPT, which reported density in the range of 0.1-0.13 pg/μm^3^ for HeLa cells.^13,42^ Importantly, when we compare the coefficient of variation (CV) between the different biophysical parameters, the CV of density are up to 10-fold smaller compared to the CV of cellular volume or mass (Figure 2G). The tight-bound density of each cell type allows us to distinguish cells based on density measurement, a property that was instrumental in cell separation with density gradient centrifugation.^8,9^ Importantly, with iDI-rSRS, we can segment the cytoplasm and nucleus to quantify their respective mass, volumes, and densities (Fig. S2). Both the cytoplasmic and nucleic drymass densities are also tightly bound with similar level of CV as the cell density, further suggesting cell-autonomous regulation of density and its importance in cell function.^7^ We note that cell density is slightly lower in nuclei compared to cytoplasm in the three cell lines we have studied, confirming recent reports of less dense cell nucleic compared to cell cytoplasm despite the large amount of DNA, histones, and other proteins packed into the finite space.^25,42^ Interestingly, the ratio between nucleic to cytoplasm density is also different based on cell types: A549 has a higher ratio than HEK and HeLa cells. This finding suggests that the dry mass density in nuclei and cytoplasm is robustly controlled, but likely by different yet unknown mechanism. Nevertheless, those differences in cell density and organelle density demonstrate the potential for cell type differentiation.

It is known that intracellular density can change in response to environmental perturbation. One advantage of iDI-rSRS over both SMR and QPT for density measurement is that iDI-rSRS provides fast and real-time density readout without the need to change medium or 3D reconstruction. We tested the sensitivity of our iDI-rSRS technique to osmotic challenges. First, we imaged A549 cells grown in an on-stage incubator with control growth media (300□mOsmol) to verify that our laser powers (30 and 35 mW for pump and Stokes, respectively) would not perturb our cells. As expected, without perturbation of the extracellular osmolarity, the cells displayed no change in density for the duration of time-lapse imaging (Fig 3A). The observed fluctuation of drymass density is 0.18 fg/μm^3^, which reflects measurement precision due to random fluctuations in laser and minute movement of the sample. We then modulated the osmolarity of the growth medium through the flow cell. Cells are imaged every 10 s for 13 mins following osmotic challenges. Introduction of hypertonic medium (+Δ200 mOsmol) leads to a quick increase in the density of A549 cells (solid line, 34□±□4% at 300 s), indicating that water exited the cells (Fig 3A, B). The density reached its peak at around 3-4 mins, and then gradually returned to its original values. This is due to a programmed response to increased osmotic pressure as regulatory volume decrease causes cells to release ions to counteract osmolarity change.^43^ Conversely, when the hypotonic medium (−Δ150□mOsmol) was introduced the change in density (solid line, −26□±2% at 300 s) was in the opposite direction. Again, the cells reached its peak change at 3-4 mins and then recovered close to their original density values as the regulatory volume increase triggers the influx of osmolytes.

**Figure 3.**
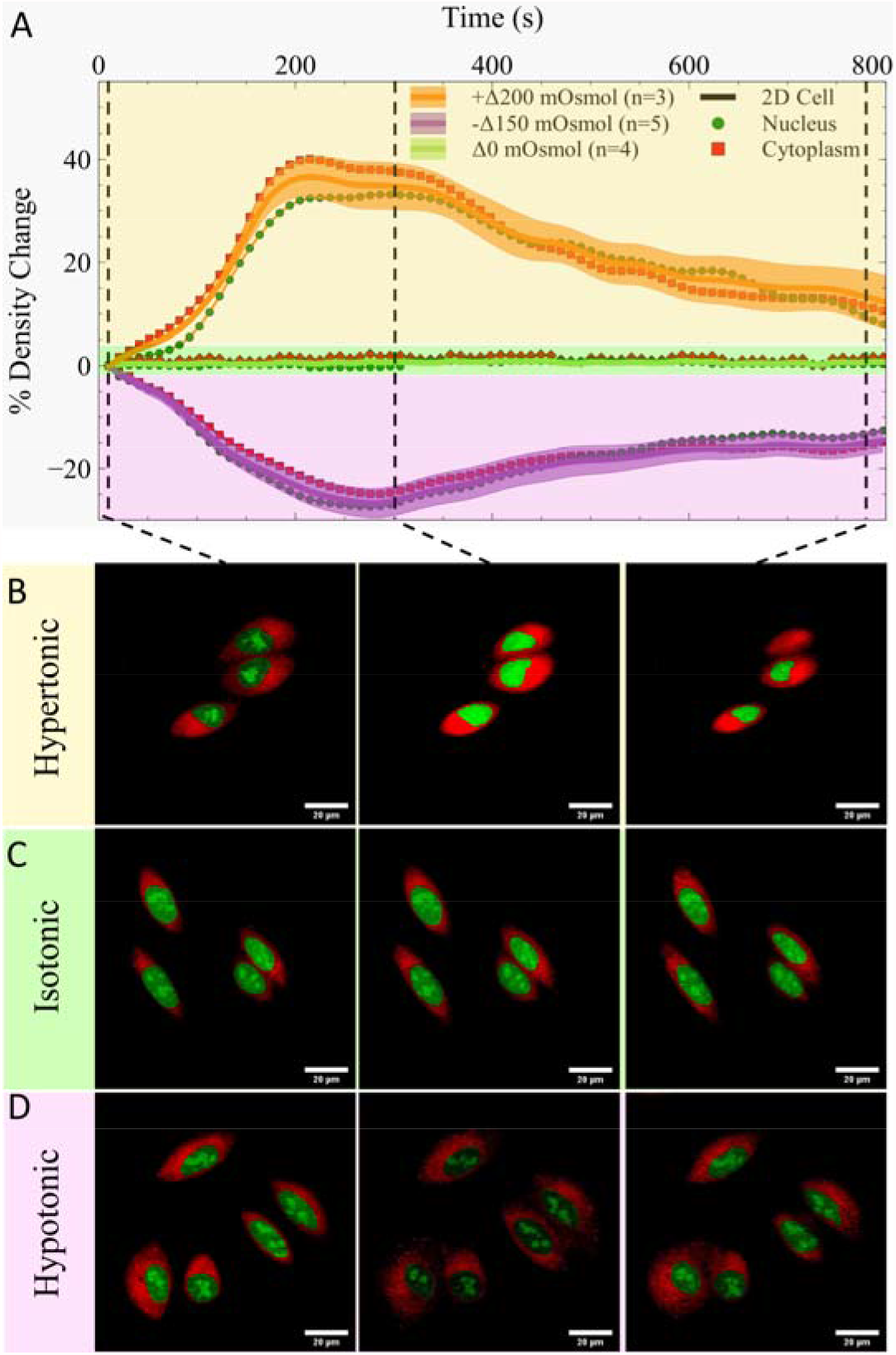
A) Relative change in density time traces for A549 cells subjected to osmotic perturbations. Representative images of: A) hypertonic medium (+Δ200 mOsmol); B) isotonic medium (Δ0 mOsmol); and C) hypotonic medium (-Δ150 mOsmol) at 0, 300, and 800 s time marks as indicated by the dotted lines. Nuclei and cytoplasm are shown in green and red color, respectively.

We further segmented the cell into cytoplasmic and nucleic regions to discern the effects, if any, of osmotic stress on these subcellular compartments. Both the nucleic and cytoplasmic densities changed in similar fashion, while maintaining the ratio between the two. It suggests that relative mass densities of cellular compartments are robustly conserved in the cell, hinting at underlying functional importance. However, the mechanism of such regulation is unknown. Nevertheless, cells can alter gene expression and metabolic activity in response to changes in osmotic environment.^44^ Thus, osmotic stress is a potent regulator of normal cell function and iDI-rSRS offers a convenient approach to monitor osmotic response over time for further investigation of its functional importance. We note that the time lapse measurements are done at a single SRS imaging plane to increase the imaging speed and decrease photodamage and thermal perturbation to the cell. To verify that the density mapping at a single plane is representative of the whole cells, we also imaged the same cells in 3D at two time points: 0 and 300 s. From our 3D density measurements (total mass divided by volume), the relative peak density change for a hypertonic and hypotonic solution are 37% and −27%, respectively. From our 3D measurements, the relative density change for a hypertonic and hypotonic solution are 37% and −31%, respectively. From these results, we can observe that our osmotic challenge results in 2D and 3D agree relatively well with each other with the small discrepancy due to differences in the contribution of nuclei to the average density as well as potential segmentation errors in 3D.

While the osmotic response and its mechanism of cells are well studied, less is known about the self-directed modulation of intracellular density in physiological conditions. Here we applied our iDI-rSRS method to quantify cell density at three different cell states: interphase, mitosis, and apoptosis. To detect mitotic cells, we incubated live A549 cells with Hoescht 33342 dye, which stains the cellular DNA, allowing for widefield epi-fluorescence detection of highly compacted DNA during the prometaphase/metaphase stage of mitosis (Fig 4C). During mitosis, adherent cells round up to facilitate accurate chromosome segregation and division.^45^ Using iDI-rSRS, we quantify and differentiate the intracellular mass densities of interphase (control) and mitotic cells as 0.103 ± 0.0056 pg/μm^3^ and 0.0958 ± 0.0015 pg/μm^3^, respectively. Mitotic swelling has been observed before.^12,13^ A fast and transient volume increase of up to 20% has been linked to the mitotic state, which explains the density decrease we observed. From the change in density, we can estimate that the average A549 cell volume expands ∼7.5% during mitosis.

**Figure 4.**
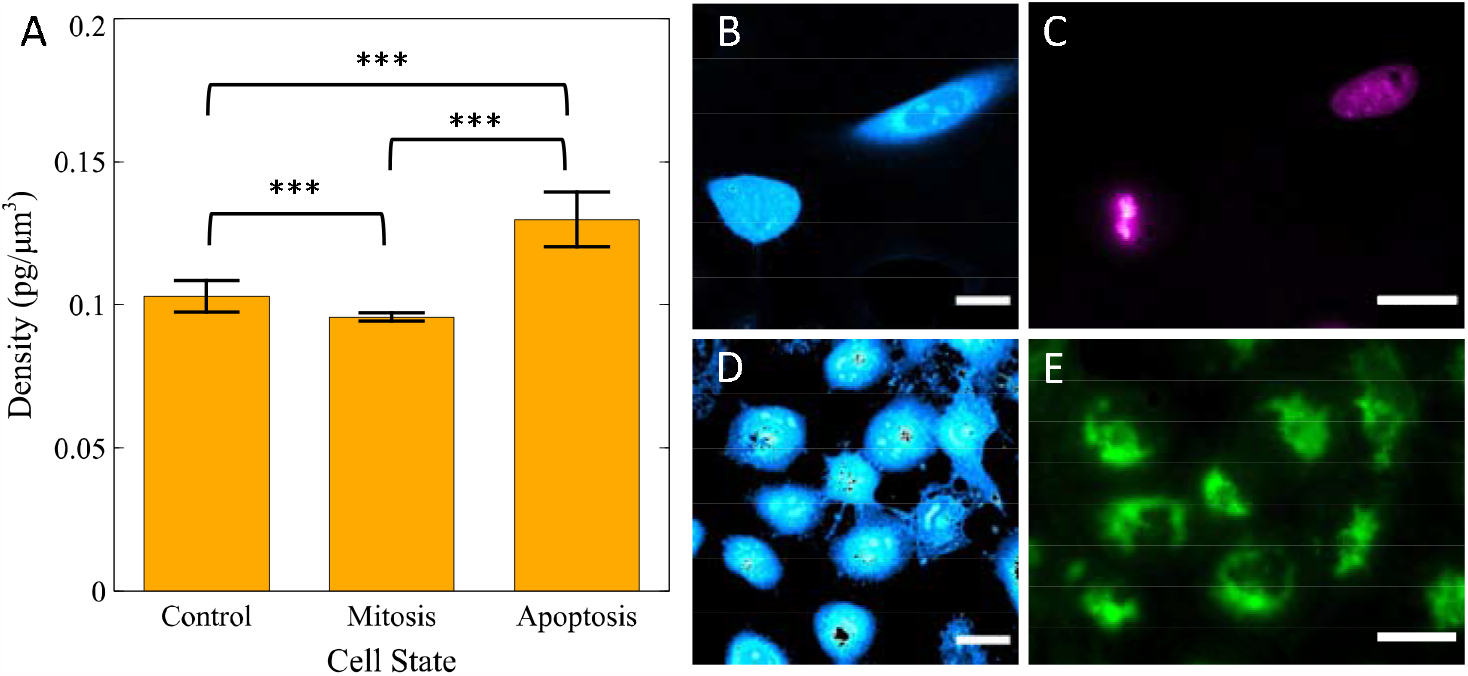
A) Density measurements of interphase (n = 27), mitotic (n = 12), and apoptotic (n = 40) A549 cells. B) Representative SRS image of A549 cells with a mitotic cell detected by C) Hoescht 33342 staining. D) Representative SRS image of apoptotic A549 cells induced by 2 mM H_2_O_2_ and detected by E) Annexin V staining. *** p≤0.001

Next, we measured the intracellular density of apoptotic cells induced by hydrogen peroxide (H_2_O_2_). H_2_O_2_ is cytotoxic and causes apoptosis at a wide concentration range. To detect apoptotic cells, we incubated live A549 cells with Annexin V, which has a strong, Ca^2+^-dependent affinity for phosphatidylserine residues that appear on the surface of the plasma membrane of apoptotic cells (Fig. 4E). We measured the apoptotic cells with an intracellular density of 0.129 ± 0.0096 pg/um^3^. A major hallmark of apoptosis is normotonic shrinkage of cells. Apoptotic volume decrease, a consequence of K^+^ release from cells, has been shown to be a prerequisite of apoptosis.^46,47^ Large increase in intracellular density, as measured here, is a direct reflection of cell condensation. Based on density measurements of A549 cells, the volume of apoptotic cells decreases ∼25% on average. The large range of density observed is because cells are at different stages of apoptosis. Nevertheless, because of the tight range of density regulation, we can effectively use iDI-rSRS measured density to differentiate cell states. Additionally, the morphological features of apoptotic and mitotic cells are also distinct as nuclei with lower density is no longer visible, further facilitating label-free identification of those cells.

Lastly, we demonstrate that our iDI-rSRS method is capable of imaging cells in a multicellular environment, which is a unique advantage over existing density imaging techniques (Figure 5A). Like other high resolution optical imaging techniques, SRS is susceptible to aberration and light scattering. Although cell density is roughly constant throughout the spheroid, the protein SRS signal decreases with imaging depth due to increased light scattering (reduces excitation laser intensity) and sample induced aberration (reduce excitation focal quality).^48^ Furthermore, closer to the boundary of the coverslip (z = 0) SRS signal also deteriorates because the SRS excitation volume extends into the coverslip. If not corrected, the obtained drymass density using SRS intensity calibration alone leads to erroneous results and shows an obvious depth dependence (Figure. 5B). With iDI-rSRS imaging, those changes associated with excitation volume and intensity cancel out for the two SRS channels. Correct drymass density can be readily obtained using the same calibration curve obtained earlier (Figure 5B). As expected, we observe very little density variation as we image through the entire tumor spheroid. Near the top of the spheroid, density decreases because the focal volume only contains partial cell which leads to underestimation. Most importantly, we observed that the same cells cultured as 2D monolayer showed a much lower intracellular density than those cultured in a 3D spheroid, particularly the nuclei density. It has been widely recognized that cells grown in 3D exhibit vastly different gene expression, signaling, and metabolic profiles from cells grown in monolayers on a rigid substrate.^49,50^ The observed density difference could be due to biomechanical differences between 2D and 3D cell environment. Such a difference directly impacts macromolecular crowding in cell nuclei. Consequently, gene transcription and DNA replication, processes that uses macromolecular machines, are directly affected because of changes in concentration and diffusion.^51,52^ Increased nucleic density in 3D cell cultures thus has far reaching consequences in the physiological function of the cell through modulation of gene activities. To the best of our knowledge, this is the first direct observation of intracellular density difference between 2D and 3D cell cultures, which further highlight the advantage of using 3D cell culture to better mimic physiological function of cells *in vivo*. The demonstrated *in situ* density imaging capability in multicellular system is an important step towards a better understanding of density regulation in tissue and its functional importance.

**Figure 5.**
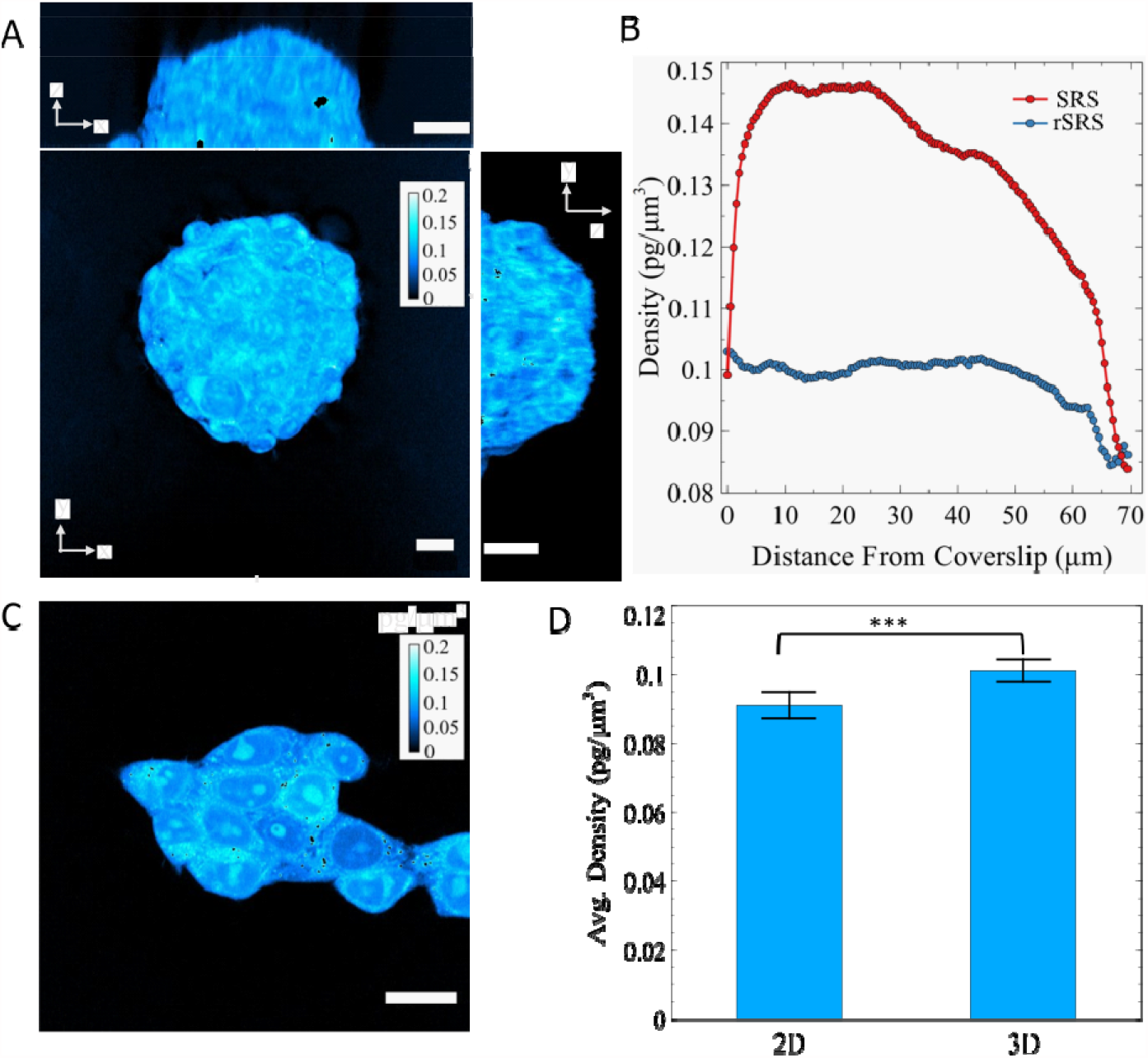
Ratiometric SRS enables robust quantification by eliminating undesirable intensity variation. A) Ratiometric 3D imaging of a 70 nm HEK293 spheroid. B) Cross-sectional density as a function of distance from the bottom to the top of the spheroid. C) Ratiometric 2D image of HEK293 cells D) Density comparison between HEK293 cells in 2D monolayer (n = 54) and 3D spheroid (n = 32). *** p≤0.001

## Conclusion

The regulation of volume and drymass in cells is essential for maintaining tissue homeostasis. Cells adjust their density during important processes such as differentiation, metabolic switch, disease state, mitosis, and apoptosis. The functional importance of density change during these processes is mostly unknown. A major analytical challenge is accurate monitoring of density changes of individual cells within their native environment. Existing methods for measuring intracellular density are either indirect measurement of surrogate parameters (e.g., refractive index) or require specific sample preparations which limits their applicability. Here, we developed a novel iDI-rSRS technique to measure the intracellular mass density of single-cells in 2D and 3D models *in situ* with subcellular resolution. The measurement principle is based on converting SRS intensity to cell drymass density and using water as an internal standard to correct for depth-dependent light scattering and aberration. Using our recently developed broadband laser and an orthogonal modulation technique, ratiometric SRS imaging at 2930 and 3170 cm^-1^ were simultaneously captured. The resulting intensity ratio was calculated to determine drymass density after proper calibration with QPM. We obtained an average accuracy of approximately 0.004 pg/μm^3^.

Using iDI-rSRS, we showed that cells tightly regulate their drymass densities, much more than their mass or volume. This tight control is present both at the cell level and the nucleus level, indicating that maintenance of cell drymass density is important for cell function. Indeed, cellular state changes such as mitosis and apoptosis are accompanied by well-controlled changes in cell density. We observed ∼7.5% intracellular density decreases and ∼25% increase in mitotic and apoptotic cell, respectively, which are consistent with previous reports. With iDI-rSRS microscopy, however, we demonstrate an appealing alternative for screening and examining cell states at a higher experimental throughput than QPT and SMR. This capability is further demonstrated in dynamic imaging of osmotic response of live cells. We showed that in response to hypoosmotic and hyperosmotic challenge, cells swell and shrink, respectively, and reach maximum density change at ∼3 mins. Cells then slowly adjust their density back towards their original density. We found that both cytoplasmic density and nucleic density change similarly, suggesting that the relative density ratio between nuclei and cytoplasm is maintained under perturbation.

Because intracellular density influences macromolecular concentration and crowding, it can directly impact gene activity through modulation of diffusion rates of transcription factors and mRNA.^5,51,53^ Crowding is also crucial for the efficient function of biological systems by promoting entropically favorable molecular association events. In the cytoplasm, crowding has been shown to induce phase separation with biological condensates that may play important roles in diverse fields including cell division, development, cancer, and neurodegenerative disease.^54^ However, tools for measuring crowding is rather limited.^55^ iDI-rSRS provides a powerful alternative to existing tools due to its high spatial resolution and high throughput. Importantly, it is directly applicable to multicellular systems. We showed that in a 3D multicellular spheroid model, cells have higher density than cells grown in a monolayer. The large change in nucleic density implies that gene activity will alter significantly, which contributes to the long observed phenotypic differences between 2D and 3D cell culture.^56^ Notably, the real-time imaging capability of iRI-rSRS makes it particularly useful for the characterization of dynamic systems. Coupled with an epi-detection SRS microscope, iDI-rSRS imaging can potentially enable *in vivo* intracellular density measurements in live animals. Combining iDI-rSRS with deep learning can be used to improve signal-to-noise ratios and measurement accuracy *in vivo*.^57^

It is important to point out that iRI-rSRS accuracy can be influenced by a couple of factors. The most important one being cellular composition. Because we only image at one Raman transition that corresponds to the protein peak and rely on the calibration of correction factor that converts rSRS intensity to drymass density using QPM drymass measurement, absolute accuracy of drymass density depends on QPM drymass accuracy and local variations of cellular composition. QPM drymass measurement is a relatively well-established technique, but its accuracy will be affected by the overall compositional variation of cells as well.^58^ Such effect still need to be explored. Within the cell, at different cellular compartments or individual imaging voxels, the composition of proteins, lipids, nucleic acids, and carbohydrate will have much large variations than cell to cell variation. iRI-rSRS does not take that into account. QPT face the same challenge. To convert the obtained RI into drymass density requires the specific refractive index increment, which depends on macromolecular composition that is typically unknown. For iRI-rSRS, we indirectly estimated the error by comparing rSRS drymass (assuming all SRS signals are from proteins) with QPM drymass to be ∼2%. Nonetheless, absolute density values at high resolution should be used with caution. The overall density change of a cell and cell nuclei is more accurate because compositional change of a single cell is likely small. It is also possible to perform SRS imaging at multiple Raman bands to further separate contribution of proteins and lipids, at the expense of increased computational complexity and imaging throughput.^59^ The second complicating factor is lipid droplets. Lipid droplets are dynamic organelles that stores neutral lipids. Their density is very high and is entirely different from intracellular density. The content of lipid droplets can vary greatly depending on cell type and metabolic status. Unlike global cell measurement like SMR, in iRI-rSRS experiments, we can separate out the contribution of lipid droplets before density calculation through image thresholding. The quantity of lipid droplets can be easily determined, providing additional information about cell physiology. Our iDI-rSRS technique is complementary to existing SMR technique and compares favorably to QPT for intracellular density imaging.^60^ Its versatility in working with multicellular systems and fast imaging capability for dynamic studies is a valuable addition to the existing toolset for studying the physiological basis and functional importance of density regulation and macromolecular crowding.

