## Supplemental file for "Quantitative imaging of intracellular density with ratiometric stimulated Raman scattering microscopy"

**Optical Quantification of Intracellular Density via Ratiometric Stimulated Raman Scattering Microscopy**

**Table of contents**

- Drymass density calibration for rSRS and error estimation
- **Figure S1:** DMSO and Methanol Spectral Calibration
- **Figure S2:** Nuclei and Cytoplasm Density Comparison Between 3 Different Cell Types
- **Figure S3:** 3D imaging of A549 cells

**Drymass density calibration of rSRS and error estimation**

We first derive the relationship between SRS intensity and drymass without considering signal normalization by water. SRS signal at 2930 cm^-1^ has three main contributions – proteins, lipids, and nucleic acids. For each voxel in the cell, assume that the drymass density is *D* (drymass *M* divided by volume *V*) and the mass fraction of protein, lipids, nucleic acids, and sugars are *α_p_*, *α_l,_* *α_n_*, and *α_s_* respectively. For each species, SRS intensity is linearly proportional to its dry mass density. The proportional constant *R* can be determined from calibration of pure species as shown in Figure 1B (in the manuscript we used BSA to represent proteins to obtain *R_p_*).

$$I=R_{p}D_{p}+R_{l}D_{l}+R_{n}D_{n}+R_{s}D_{s}$$

$={(\alpha}_{p}R_{p}+\alpha_{l}R_{l}+\alpha_{n}R_{n}+\alpha_{s}R_{s})D$	(1)

Total SRS signal *I* of the cell will then be a sum of the contribution of all four species integrated over the cell volume.

$$\int IdV=\int{(\alpha}_{p}R_{p}+\alpha_{l}R_{l}+\alpha_{n}R_{n}+\alpha_{s}R_{s})DdV$$

$={(\alpha}_{p}R_{p}+\alpha_{l}R_{l}+\alpha_{n}R_{n}+\alpha_{s}R_{s})M$	(2)

Therefore, the total SRS intensity (integrated over the cell volume) is linearly proportional to drymass *M* measured with QPM. As a result, we can calculate the coefficient ${(\alpha}_{p}R_{p}+\alpha_{l}R_{l}+\alpha_{n}R_{n}+\alpha_{s}R_{s})$ that converts SRS intensity to drymass density by comparing SRS and QPM measurements directly. Throughout the manuscript, we used this calibrated coefficient to determine drymass density for all the cells. We note that at each imaging voxel the drymass composition of biomolecules will be different, which leads to errors in absolute quantification at the individual voxel level. Quantitative phase tomography (QPT) has a similar problem because the refractive index increment depends on drymass composition. However, the average density of a cell measured by SRS will still be accurate. Relative changes of density (such as shown in the osmotic challenge experiment) is also accurate because the drymass composition does not change rapidly.

The use of rSRS does not change the aforementioned equations if we simply replace the SRS intensity *I* and proportional constant *R* with the corresponding numbers from rSRS measurements. The use of ratiometric measurements normalizes signal change due to aberration and scattering. It is a more robust measurement compare to intensity.

Alternatively, we can calculate approximate SRS drymass *M’* based on solution calibration measurements if we assume all drymass molecules have the same *R* as protein

$\int IdV=R_{p}M'$	(3)

Substituting into equation (2)

$M^{'}=\frac{R_{p}}{{(\alpha}_{p}R_{p}+\alpha_{l}R_{l}+\alpha_{n}R_{n}+\alpha_{s}R_{s})}M= \gamma M$	(4)

As a result, we can calculate cell drymass directly from measured SRS drymass with a correction factor *γ* (Figure 1B).

Next, we estimate the drymass density error caused by variations in drymass composition. We can assess this error by comparing directly SRS measured drymass *M’* (before correction) and QPM drymass *M*. When comparing *M’* with drymass measured by QPM, a linear relationship with a slope of 1.018 (which is the correct factor) is obtained. The small difference of these two measurements (<2%) indicates that the drymass approximation is reasonably accurate even though we do not distinguish proteins from other biomolecules in SRS. This is largely because protein accounts for the majority of the drymass and the contribution of other species to SRS at the protein Raman peak further reduce the error with the approximation. Therefore, the intracellular density error associated with drymass composition variation in the cell should be small.

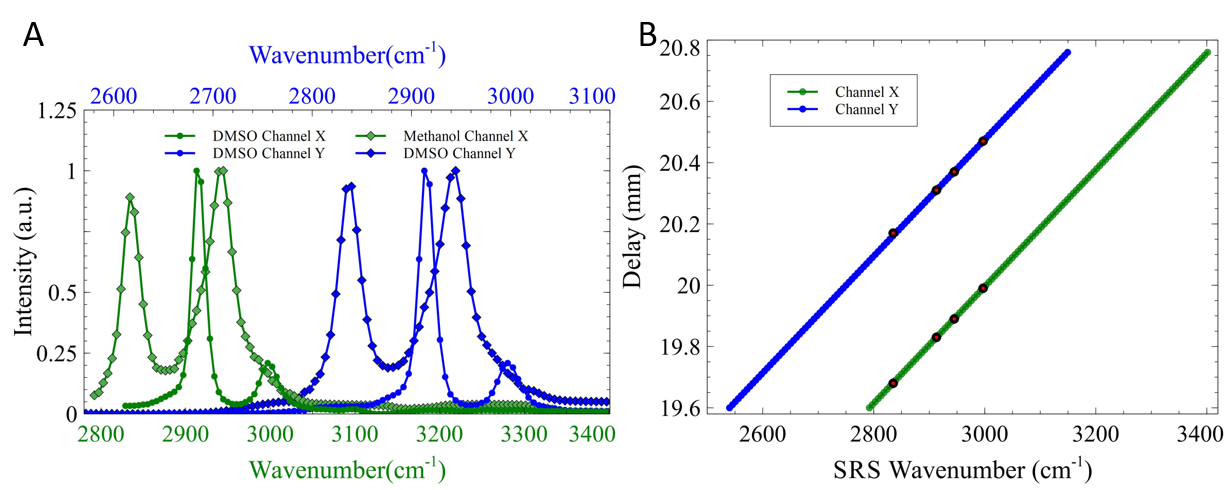

Figure S1: Calibration of SRS imaging setup based on a parabolically amplified femtosecond dual beam laser system. A) Simultaneous collection of 2-channel SRS spectra arising from CH stretching vibrations of DMSO and Methanol. B) Calibration of SRS wavenumber with respect to pump-Stokes pulse delay using Raman spectra of DMSO and Methanol.

Figure S2. Average A) mass, B) volume, C) density between the three different cell types along with their cytoplasmic and nucleic contributions. Error bars are ± one standard deviation of the mean.

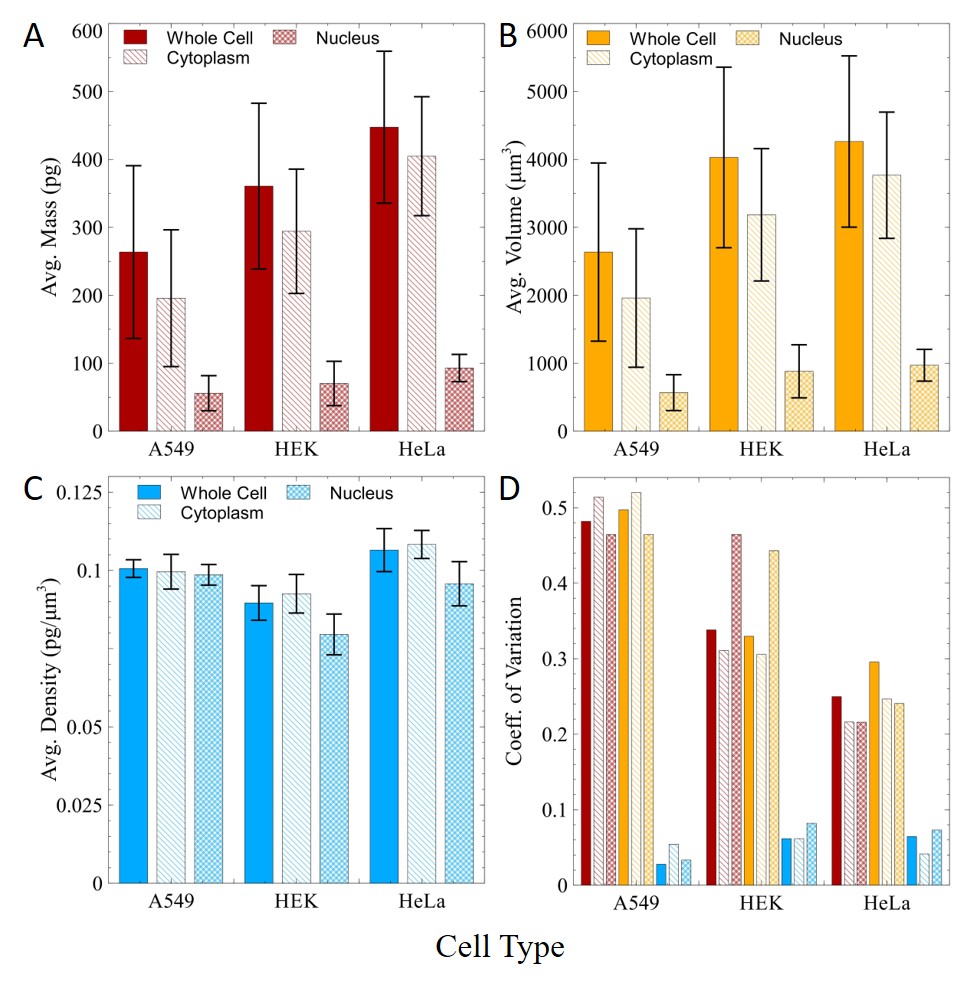

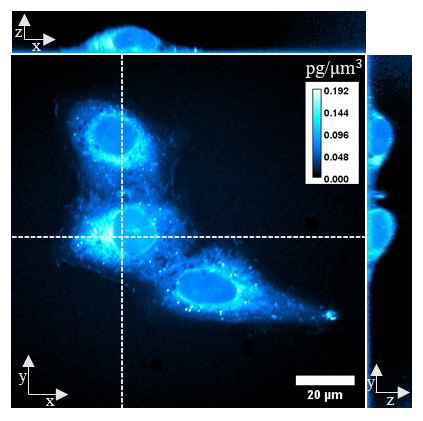

Figure S3. 3D Imaging showing the intracellular density distribution of A549 cells.
